# Studying bone marrow-derived macrophages needs to consider the influence of cells apart from macrophage, especially the selection of markers

**DOI:** 10.1101/2020.10.01.305748

**Authors:** Chenyu Chu, Shengan Rung, Yufei Wang, Yili Qu, Yi Man

## Abstract

Culturing macrophage in vitro is an important means to understand its reaction towards co-culture substances. However, the source of macrophages can be either purchased from specific cell line or extracted from bone marrow as differentiated macrophages. In order to assure the precision, when adopting extracted primary cell, screening in advance will be prominent before any processes to avoid results reserving that of non-macrophages. Here, we perform single-cell sequencing on open wound of skin in mice and focusing on the secreted proteins and surface markers related to traditional macrophage phenotypes (M1/M2) to ensure the importance of screening. The expression of CAMP when macrophages fight against candida albicans is another target to see its relationship with current classification. And results showed that identifying its phenotype without screening macrophages will far from the exact situation, and the expression of CAMP cannot be carried out by the traditional M1 and M2 macrophage. Thus, determining phenotype of macrophages based on function would be a promising way.

## Introduction

Macrophage is one of the most important defenders and regulators of the immune response, which will be activated by the immunological recognition of pathogens and foreign bodies. The timely shifting from M1 towards the M2 phenotype of Macrophage is usually concerned with tissue repair. In contrast, Gao et al. mentioned that an opposite strategy was needed for tackling Candida albicans with the demand effectively to improve the survival rate after being infected ^1^. To evaluate the nano-trinity treatment effect on phenotypes of macrophage and its expression, flow, western blot (WB), and ELISA kits were adopted. However, the controversies in this study not only appeared in the selected markers and the grouping process which seem lacking comprehensive consideration, but also emerged in the glancing description to the interaction between specific phenotypes of macrophages and Candida. In this study, we discussed the related topic with single cell RNA sequencing data, investigated the contradictory results and analysis the underlying way of classification of phenotype during biomaterials mediated host defense against Candida albicans.

## Method and materials

The original data is obtained from previous database2 and processed through Seurat3 (https://satijalab.org/seurat/) and Loupe Browser v4.0 (10xgenomics, USA) for subsequent analysis.

## Results

### Macrophage-specific markers should be detected before determining its phenotype

In order to confirm the specificity of the macrophage phenotypic markers (Fig. 1A, B) and proteins (Fig. 1C-G) secreted by specific phenotypes, we analyzed the expression of each cluster in open wound of skin. In addition, it was found that none of the detected markers are completely specific; most of the targeting gene have close relationship with neutrophils (Fig. 1H) or fibroblasts (Fig. 1I). That is, through those markers to determine macrophage phenotype from an obvious potentially complex environment without screening the specific marker of macrophages like F4/80 in advance, the result is prone to false positives from other non-macrophage cells.

**Figure 1.**
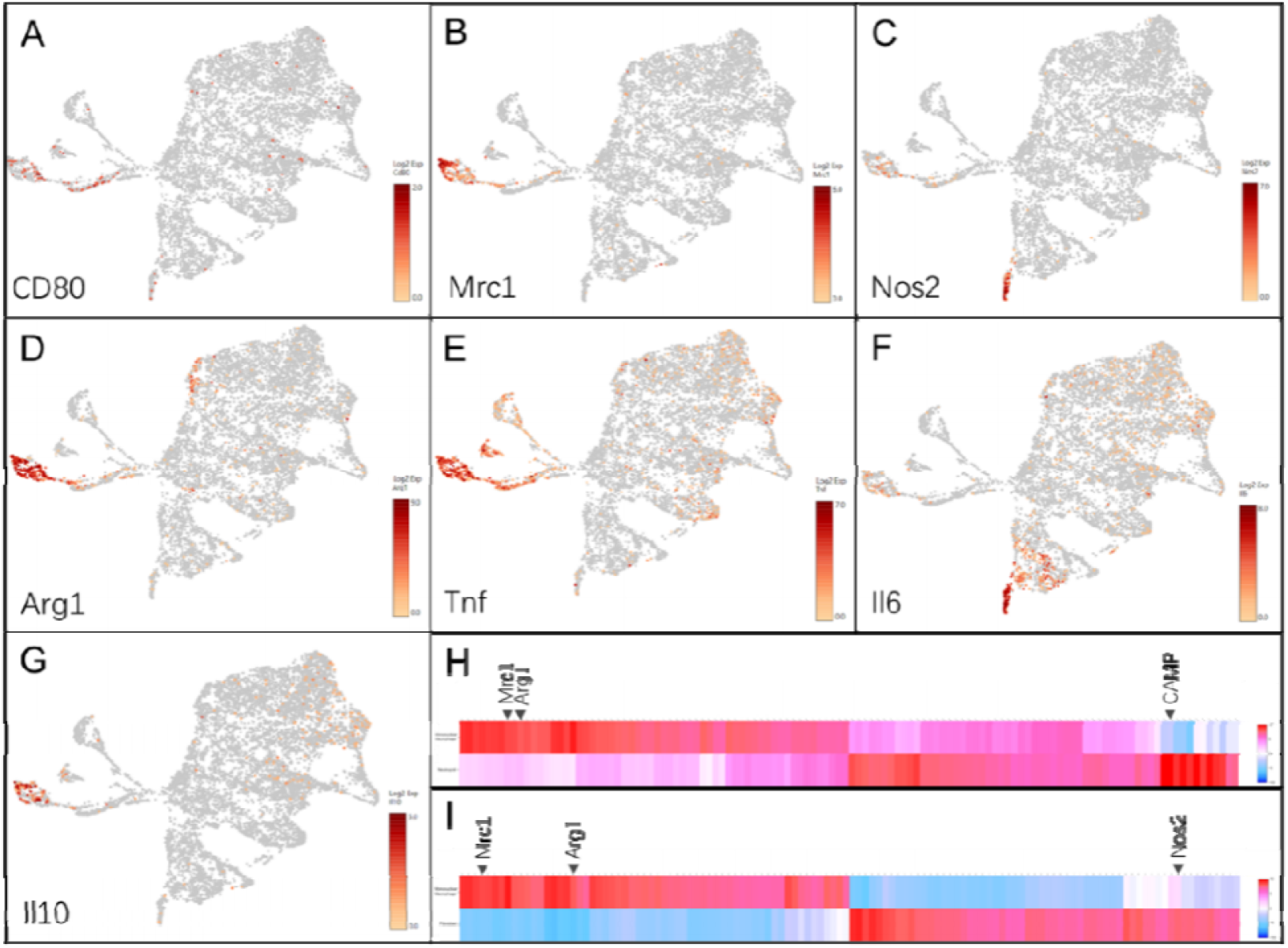
Single-cell sequencing result of skin open wound. Expression of gene (A) CD80, (B) Mrc1, (C) Nos2, (D) Arg1, (E) Tnf, (F) Il6, (G) Il10 and heat map of monomolecular macrophage versus (H) neutrophil and (I) fibroblast cluster

### The expression of Camp is independent of traditional M1/M2 phenotypic markers

Comparing the gene expression of macrophages and neutrophils, Arg1 could be expressed by both cells in open wound of skin; this phenomenon can also be seen in Nos2 (Fig. 2A, B and D). In addition, we found that the expression of Camp has nothing to do with traditional macrophage phenotypic markers (M1-Nos2/M2-Arg1), for neither Arg1+Camp+ nor Nos2+Camp+ cells exist (Fig. 2E and 2F). Moreover, Camp could only be seen in neutrophil cluster with relative low expression (Fig. 2C).

**Figure 2.**
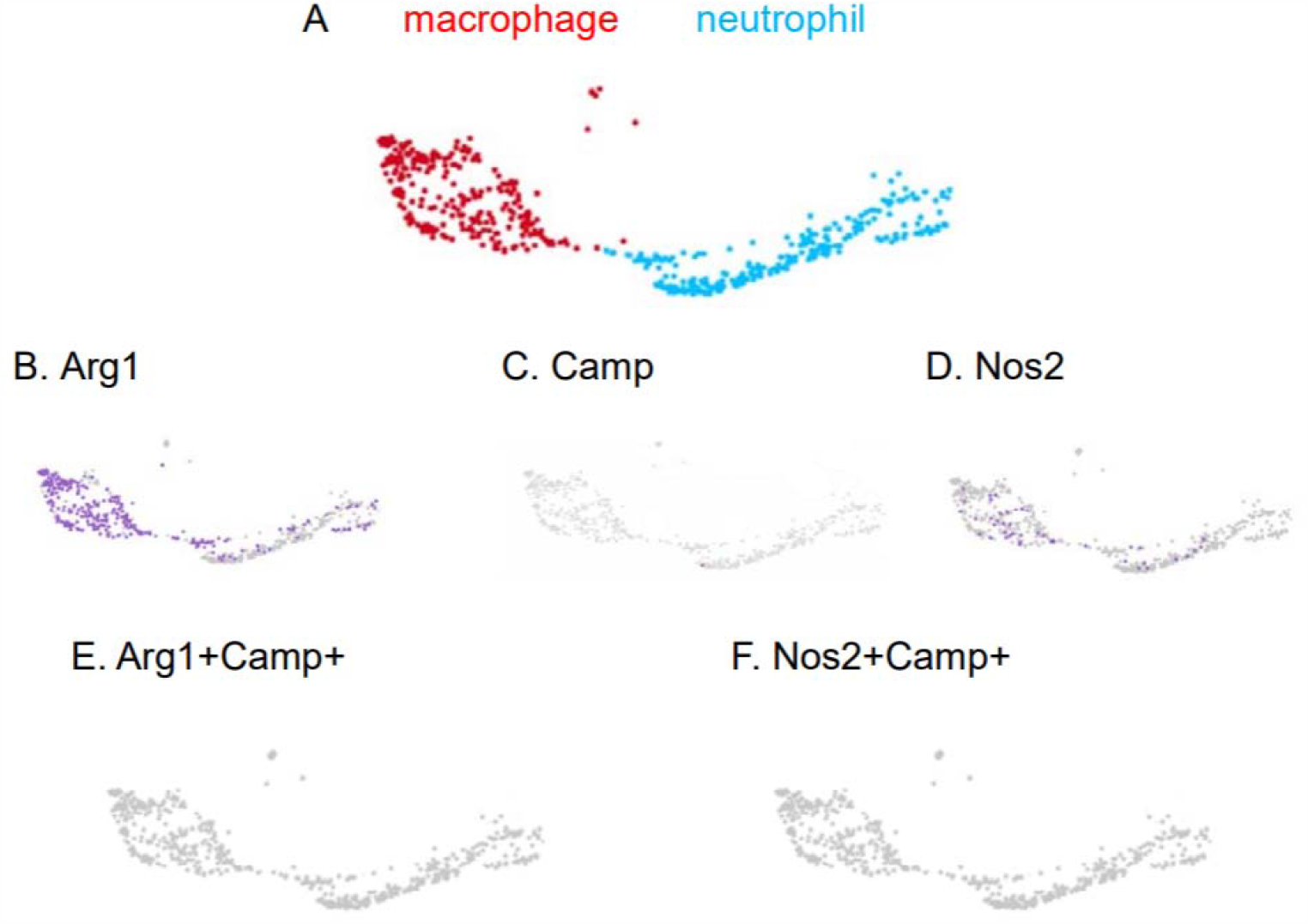
(A) Single-cell sequencing result of monomolecular macrophage (red) and neutrophil (blue). The expression of (B) Arg1, (C) Camp, (D) Nos2, (E) Arg1+Camp+ and (F) Nos2+Camp+

## Discussion

First, the phenotype of both J774A.1 macrophage cell line and bone marrow-derived macrophages (BMDMs) were detected by simultaneously incubating with F4 / 80 and CD206 (M2 phenotype) or CD80 (M1 phenotype), respectively. This indifferent treatment would accompany inevitable error to BMDMs, which would cause the data of flow cytometry included CD80^+^ and CD206^+^ of non-macrophage cells and increased false-positive results. That is, BMDMs, unlike J774A.1 macrophage, is difficult to avoid the presence of multiple types of cells apart from macrophages. As extracted *in vivo*, BMDMs is essentially different from the cell line. Moreover, merely CD80 or CD206 could not represented for a certain phenotype; more markers should be adopted. In addition, the F4 / 80 staining results showed that the positive population had no obvious clustering that was difficult to demonstrate the existence of an F4 / 80^high^ population in BMDMs. To resolve this problem, conducting flow cytometry for phenotype indication should begin with extracting CD11b^+^F4 / 80^+^ population from the entire BMDMs to determine macrophage, then analyzing those phenotypic M1/M2 specific markers. Besides, according to the previous study, macrophages with both M1 and M2 markers must also be considered, therefore it would be better to stained CD80 and CD206 at the same time based on F4 / 80^high^ population (CD80^+^CD206^-^ represented for M1 phenotype; CD80^-^CD206^+^ represented for M2 phenotype). As for, the secretion of iNOS, Arg1, TNF-α, IL-6, and IL-10 to determine macrophage polarization also neglected whether they are specific for macrophage. Especially iNOS (encoded by Nos2), which was different from Arg1, our results showed that it could be secreted by neutrophil and fibroblast ^2^. Therefore, without quality control, WB and ELISA results of BMDMs must include the expression levels of other bone marrow-derived cells. Such protein quantitative results cannot correspond to flow results. In particular, separating macrophages from BMDMs before processing any assays, could better reflect the real situation. Last but not least, Candida albicans could reduce the production of NO, not by inhibiting the expression of Nos2 mRNA, but lowering the activity of iNOS enzyme and decreasing the expression of iNOS protein ^3^. Whereas other researchers suggested that it is due to the promotion of Arg1 through chitin from candida albicans in the competition for the L-arginine substrate. With the help of Arg1 inhibitor, the synthesis of NO could be restored while the survival rate of the host being improved ^4^. It may seem that weakening macrophages polarization towards M2 phenotype when tackling such pathogens would be a promising solution, but the host response under the attack candida albicans is not that simple. Indeed, there are corresponding studies aiming at both macrophages of murine and human that have an M1-to-M2 switch after candida albicans infection ^5, 6^, but whether this polarization trend is candida’s strategy to improve its survival rate or the result of host self-protection is still unknown. In addition, as recommended by Gao et al., increasing M1 macrophages can promote the expression of NO, ROS and enhance phagocytosis to fight candida albicans ^1^, however, candida albicans may end up resulting in the death of macrophages by transferring its metabolic way on glucose that macrophage could rival it ^7^. Thus, to confirm that macrophages can reduce the damage of Candida by enhancing M1 macrophage polarization, longer-term observation is needed. Cramp, an antimicrobial peptide, is another effective weapon of the host against candida albicans, which could be produced not only by macrophages, but also by neutrophils and epithelial cells ^8^. Other than antimicrobial peptides, Cabezón, Virginia, et al. had also clarified multiple proteins acting between macrophages and Candida, which is related to the induction of candida albicans’ apoptosis, confirming macrophages contribution ^9^. However, whether the M1 and M2 macrophage phenotypes are the key to regulating Candida infection, and how the specific phenotype of macrophages prolongs the host survival time needs further study. Base on the information mentioned above, by means of functionally relevant markers may be a more reasonable option for understanding the role of macrophages in fungal infections. All in all, the removal of pathogens by macrophages is indispensable, which functions closely related to the phenotype. However, in the grouping process, it should be considered whether the selected object composed of macrophages only. If there were other types of cells, they should be excluded before starting a phenotypic assay. The gating for the baseline and selection of markers should according to the composition of the tested samples. The phenotypes and corresponding antibacterial effects also interested us a lot. It would be our pleasure to have further communication on the perspective mentioned above and the author’s considerations for experimental design.

## Funding information

National Natural Science Foundation of China, Grant/Award Number: 81870801 and 81671023.

## Statement of conflict of interest

There are no conflicts of interest related to this manuscript.

## Notes

### Competing Interest Statement

The authors have declared no competing interest.

